# Revisiting the Need for mRNA Nucleoside Modification in CAR T Cell Engineering

**DOI:** 10.1101/2025.01.21.630650

**Authors:** Nourhan Kahwaji, Niklas Kotzian, Jasmin Melissa Prinz, Yaolin Pu, Jonas Kath, Samira Picht, Anna Luisa Hiller, Charlotte Maeve Dunne, Michael Launspach, Arnd Kleyer, David Nils Simon, Dimitrios Laurin Wagner, Chantal Pichon, Gerhard Krönke, Michael Schmueck-Henneresse, Hans-Dieter Volk, Manfred Gossen, Norman Michael Drzeniek

## Abstract

mRNA-based chimeric antigen receptor (CAR)-T cells offer the promise of enhanced safety and simplified manufacturing. However, in vitro-transcribed (IVT) mRNA is known to trigger antiviral immune responses, inflammatory signaling, and apoptosis in transfected cells. To address these challenges and enable efficient IVT-mRNA expression, modified nucleosides, such as N1-methyl-pseudouridine (m1Ψ), have become the gold standard for CAR-T cell production, albeit at increased cost.

In this study, immune responses to IVT-mRNA were evaluated across five primary human cell types, including T-cells. Unexpectedly, T-cells, unlike other immune and non-immune cell types tested, exhibited no immune activation in response to unmodified mRNA. T-cell viability and cytokine secretion patterns remained unaffected, regardless of whether unmodified mRNA was delivered via lipid nanoparticles (LNPs) or electroporation.

Furthermore, CAR expression levels in T-cells were not influenced by mRNA modification with m1Ψ or 5-methoxy-uridine (5moU) nucleosides. The absence of nucleoside modifications did not compromise CAR-T cell cytotoxic potency, demonstrating that such modifications are not required for producing functional CAR-T cells.

These findings eliminate the need for nucleoside modification in T-cell mRNA, simplifying and reducing the cost of CAR-T cell manufacturing while positioning IVT-mRNA as a highly efficient and minimally invasive tool for CAR-T cell engineering.

## Introduction

Virus-transduced CAR-T-cell therapies are often associated with high cost, burdensome logistics and lingering safety concerns [1]. Novel non-malignant applications of CAR-T-cells, such as breakthrough treatment of autoimmune disease [2] and promising elimination of fibrosis and senescence [3, 4], have sparked an acute interest in mRNA-based transient CAR-T-cell products with improved safety and reduced manufacturing complexity, that facilitates scalability[5]. mRNA-CAR-T-cells are being developed against autoimmunity [6] and cancer [7] alike. Nevertheless, mRNA production remains expensive, largely due to the use of chemically modified nucleosides.

Excessive immune activation upon recognition of IVT-mRNA by pattern recognition receptors (PRR), a major roadblock in the development of mRNA therapeutics [8], has been largely overcome by mRNA nucleoside modifications, such as N1-methyl-pseudouridine (m1Ψ) in Comirnaty and Spikevax SARS-CoV2 vaccines [9], or 5-methoxy-uridine (5moU), which has shown even greater reduction of mRNA immune activation in our previous studies [10, 11].

m1Ψ has become a gold standard in the mRNA field and to the best of our knowledge, all reports of mRNA-based CAR-T-cell engineering use m1Ψ-modified mRNA [12, 13]. However, modified nucleosides amount to ∼35% of mRNA manufacturing cost at large-scale. Surprisingly little is known about the different sensitivity of different cell types toward mRNA-triggered immune activation and thus about cell type-dependent requirements for mRNA modification.

In our previous work, we extensively characterized the immune response toward mRNA with different chemical modifications in bone marrow stromal cells [10, 11]. Asking to what extent those insights could be transferred to other cell types, here we screened responses toward unmodified and nucleoside-modified (m1Ψ, 5moU) mRNA in 5 primary human cell types, including T-cells, macrophages, endothelial, epithelial and stromal cells.

Surprisingly, and in contrast to all other cell types tested, T-cells exhibited no immune response to unmodified mRNA, regardless of the transfection method. mRNAs with different nucleoside chemistries resulted in comparable expression of CD19-CAR in primary human T-cells. To confirm that the use of unmodified mRNA does not functionally impair T-cells, we demonstrated that mRNA-CAR-T cells transfected with unmodified mRNA exhibited equivalent cytotoxic activity compared to those transfected with industry-standard m1Ψ-modified CAR-mRNA.

## Methods

### Cell Culture

Blood was drawn from n≥3 consenting adult donors (Charité ethics committee approval EA4/091/19). From the mononuclear fraction obtained by density gradient centrifugation, primary T-cells and monocytes were isolated using magnetic cell sorting with beads against CD3 and CD14 respectively (Miltenyi Biotec, Germany)[11]. T-cells were cultured and activated in ImmunoCult-XF TCell Expansion Medium with recombinant IL-2 and ImmunoCult HumanCD3/CD28/CD2 TCell-Activator (Stemcell Technologies, Canada). Macrophages were differentiated within 7 days of culture in RPMI1640 containing 10% FCS,1% Penicillin/Streptomycin and 50ng/mL M-CSF (Miltenyi) [11, 14].

Bone marrow stromal cells were supplied by the BIH Center for Regenerative Therapies (BCRT) core facility “Cell Harvesting” [15]. Written informed consent was given, and approval was obtained from the Charité IRB/local ethics committee (EA2/089/20). Human umbilical vein endothelial cells were purchased from Promocell, Germany. Tubular epithelial cells were a kind gift from Prof. Babel’s group at BCRT. Non-immune cells were cultured in low-glucose DMEM containing 10% FCS, 1% Glutamax and 1% Penicillin/Streptomycin.

### mRNA Synthesis and Transfection

DNA sequences encoding enhanced green fluorescent protein (GFP) or chimeric antigen receptor (CAR; Supplemental Table 1) were cloned into the pRNA2-(A)128 vector, amplified by PCR and transcribed using the TranscriptAidT7 kit (Thermo Fischer, USA) into mRNA containing uridine, m1Ψ or 5moU (Jena Bioscience, Germany). mRNA was purified using lithium chloride precipitation, washed in 70% ethanol and stored in aqueous solution, as previously described [11].

mRNA was complexed with Lipofectamine MessengerMax (LMM; ThermoFischer) at a ratio of 1ug:2ul [10] and added to cells at 2ug mRNA per 1*10^6^ cells. Because LMM did not transfect T-cells (Figure S1), mRNA was microfluidically formulated with Genvoy T-cell lipid mix (Cytiva, USA) using setting #3 on a NanoAssemblr Spark and added to cells after 48h of activation.

T-cells or macrophages were electroporated using a 4D-Nucleofector (Lonza, Switzerland) in P3 buffer using the EH-115 or DP-148 programs respectively.

### Quantification of mRNA Expression and Cell Death

All readouts were performed 24h post-transfection. GFP fluorescent imaging was performed on a Nikon ECLIPSE Ti (Nikon Instruments, Japan).

GFP- and CAR-expression were quantified using using MACSQuantVYB (Miltenyi) or CytoflexLX (Beckmann Coulter, USA) flowcytometers. CD19-CARs were stained using anti-Myc-Tag (CAR1[16] 1:50; 9B11; CellSignalingTechnology, USA) or anti-IgG-Fcγ (CAR2[17]; 1:50; 109-605-098; Jackson ImmunoResearch, USA) antibodies. DAPI was used (1 µg/mL) for live/dead discrimination. Data was analysed in FlowJov10; a representative gating strategy is shown in Figure S2.

Caspase activity was quantified as previously described [10].

### Cytokine and Interferon Analysis

Concentration of interferons α and β were measured by ELISA and T-cell secretomes were measured using ProteomeProfilerXL Human Cytokine array (all R&D Systems, USA) from supernatants 24h post-transfection, according to the manufacturer’s instructions.

### Cytotoxicity Assay

CAR-dependent cytotoxicity was assessed in a VITAL assay, as previously described [16, 18]. Briefly, CAR-T-cells were incubated at decreasing ratios with Nalm6 (CD19+ GFP+; target) and Nalm6 (CD19-RFP+; control) cells for 8h, stained with DAPI, and ratios of surviving target and control cells were analyzed by flow cytometry. Cytotoxicity was normalized against mock-transfected, CAR-negative T-cells using the formula:

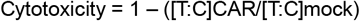

### Data Analysis

Statistical analysis was carried out in Prism10 (GraphPad Software., USA). A one-way or two-way ANOVA with Tukey’s or Dunnet’s post-hoc test was used to test for significant differences between groups. Heatmaps were generated in R and Python v3.11.0.

## Results and Discussion

Primary human T-cells, monocyte-derived macrophages, and three non-hematopoietic primary cell types representative of different tissues (human umbilical vein endothelial cells; bone marrow stromal cells; renal tubular epithelial cells) were screened for antiviral responses toward IVT-mRNA, which can reduce cell viability and impair mRNA expression [11].

Non-hematopoietic cells showed very similar interferon response, caspase activation and cell death in response to unmodified mRNA, which was greatly reduced by uridine substitution with m1Ψ and further reduced by 5moU. Macrophages showed the strongest immune response and apoptosis to unmodified mRNA (Figure1a-d; Figure S3; Figure S4). This was also reflected in the relatively large impact of uridine modification on mRNA expression in macrophages (Figure 1e-f; Figure S5).

**Figure 1.**
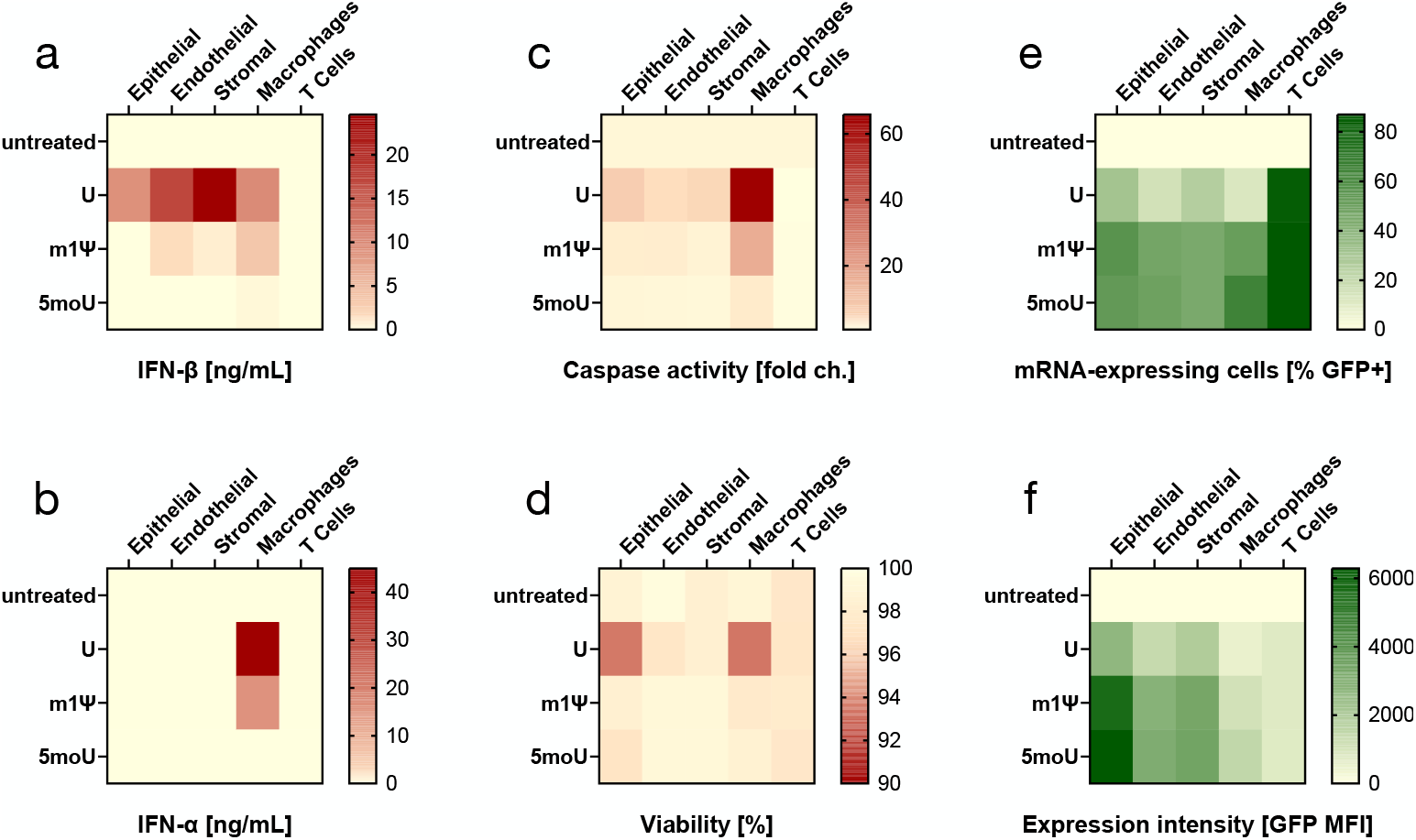
Responses toward mRNA with different nucleoside chemistry vary between cell types. Renal tubular epithelial cells, umbilical vein endothelial cells, bone marrow stromal cells, macrophages and T-cells (all primary human cells) were transfected with unmodified (U) or m1Ψ-modified or 5moU-modified GFP-encoding mRNA and cellular responses were measured 24h post-transfection: **a, b)** Secretion of type-I interferons β and α in supernatants, which are hallmarks of cellular immune responses against RNA viruses. **c)** Increase in activity of caspases 3 and 7, as an indicator of apoptosis induction. **d)** Cell viability measured using flow cytometry. The fraction of mRNA-expressing cells **(e)** and expression level **(f)** measured by quantifying GFP fluorescence using flow cytometry. n= 3-12, detailed graphs and statistical information can be found in Supplemental Figures S3-5.

Surprisingly and in contrast to all other cell types, T-cells did not show any adverse response to unmodified mRNA in terms of interferon secretion, caspase activation or cell death. Also, all three mRNAs were expressed equally, suggesting no beneficial impact of uridine modification in T-cells (Figure 1; Figure S3-5).

Except for T-cells requiring mRNA to be encapsulated in lipid nanoparticles (LNPs; Figure S1), all other cells could be transfected with lipofectamine-complexed mRNA. We therefore asked whether the lack of a recognizable immune reaction toward unmodified mRNA in T-cells depended on the LNP-based delivery.

This was not the case, as shown in side-by-side comparison of mRNA delivery using LNP versus electroporation (EP): T-cells displayed no increased cell death in response to unmodified mRNA for either transfection method (Figure 2a; Figure S6). Electroporated T-cells exhibited low viability loss in response to EP-related stress (mock-EP), which did not increase when mRNA was added.

**Figure 2.**
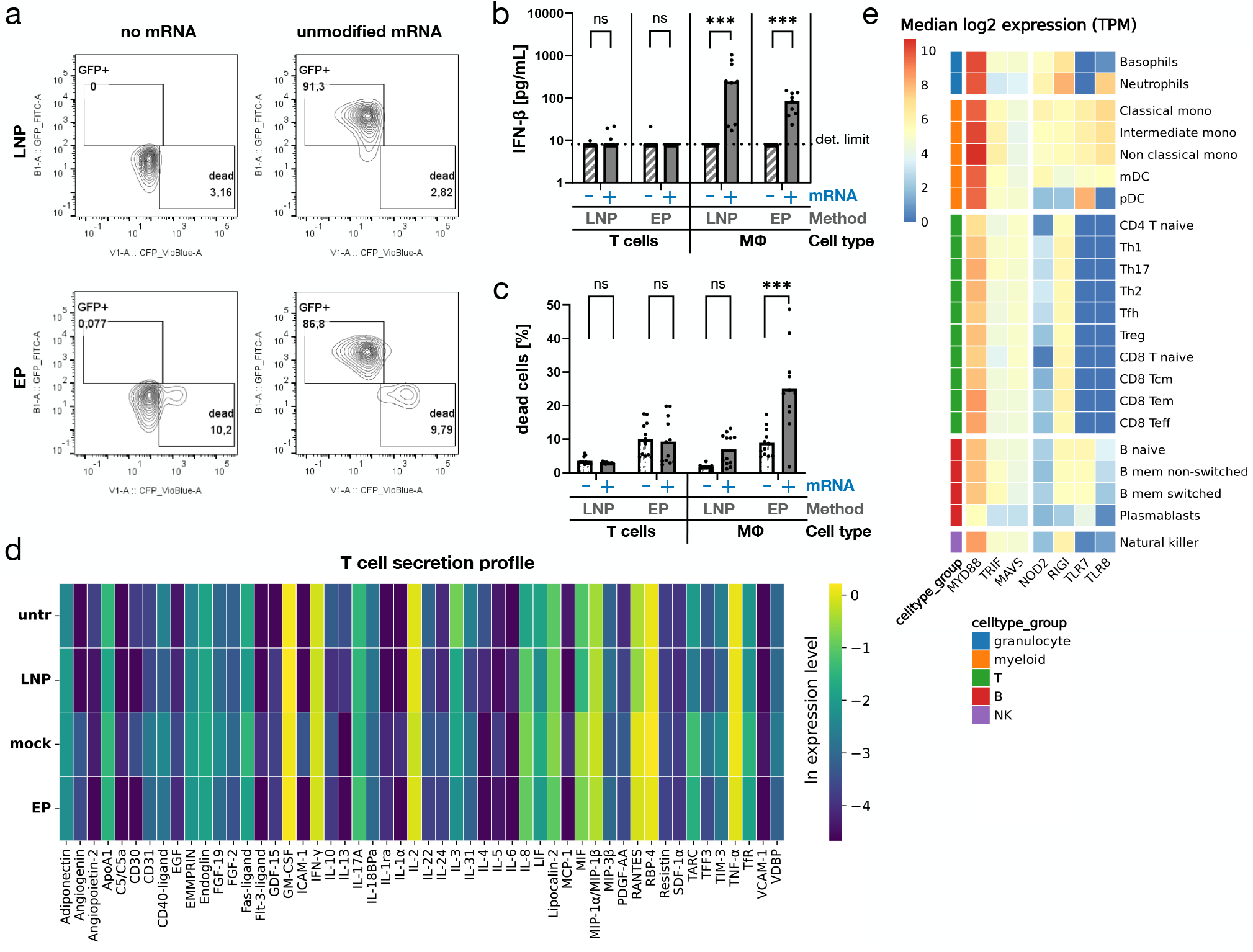
Human T-cells show no immune activation by unmodified IVT-mRNA. Unmodified mRNA was introduced into primary human T-cells using lipofection (LNP) and electroporation (EP) to exclude dependency of immune activation on the delivery mechanism. **a)** 24h following either transfection method, T-cells showed strong expression of mRNA-encoded GFP and no reduction in cell viability was observed compared to untreated or mock-electroporated cells (EP without mRNA). n=1 representative set of flow cytometry plots is shown out of n=12 from 4 biological donors (Figure S6). **b)** Interferon β secretion from T-cells was quantified using ELISA and revealed no interferon induction after transfection with unmodified mRNA through either EP or LNP. Monocyte-derived macrophages (MΦ) were used as a positive control. n=3 donors, median is shown. **c)** EP led to more T cell death compared to LNP, but in contrast to macrophages, cell death was not increased by mRNA treatment. n=4 donors. Mean is shown. **d)** A protein array measuring 105 cytokines in T cell supernatants 24h post-transfection did not reveal any secretion pattern shift upon transfection with unmodified mRNA. n=1. **e)** Publicly available bulk RNA sequencing data [19] was interrogated for transcripts of pattern recognition receptors relevant for intracellular recognition of mRNA (NOD2, RIGI, TLR7, TLR8) and related adapter proteins (MyD88, TRIF, MAVS). The analysis revealed that the two uridine-recognizing RNA receptors (TLR7, TLR8) were not expressed in any T lymphocyte subsets and their adapters MyD88 and TRIF were expressed at lower levels than in other leukocytes. TPM: transcripts per million. n=4. * p<0,05; ** p<0,01; *** p<0,001; ns= not significant.

In direct comparison with macrophages, T-cells did not show any interferon-β induction or apoptosis following transfection. Macrophages displayed both responses, although the same LNP-formulation was used to transfect both cell types (Figure 2b-c).

To assess whether mRNA transfection influenced the production of cytokines beyond type-I interferons, a protein array profiling 105 cytokines was performed on T-cell supernatants. In contrast to bespoke secretome shifts previously described in stromal cells [11], T-cells transfected with unmodified mRNA via either EP or LNP exhibited indistinguishable cytokine secretion patterns (Figure 2d). Notably, the production of key T-cell cytokines, including IFN-γ, IL-2 or the inflammation marker TNFα, remained unaffected by mRNA treatment.

In conclusion, multidimensional profiling of primary human T-cells revealed no evidence of immune activation or downstream side effects in response to unmodified mRNA.

Interrogations of RNA sequencing data from two independent sources [19, 20] revealed that among RNA-sensing PRRs, TLRs 7 and 8, which recognize unmodified uridine were expressed in T-cells at lower levels than in other leukocytes, suggesting that T-cells might not be equipped to recognize uridine-containing mRNA as a pathogen-associated pattern (Figure 2e, Figure S7).

While an in-depth characterization of RNA-sensing pathways in T-cells was outside the scope of this study, our aim was to determine whether the costly nucleoside modifications traditionally used in mRNA-based CAR-T cell production are necessary for generating potent CAR-T cells (Figure 3a). Unmodified, m1Ψ-modified, and 5moU-modified CAR-mRNA resulted in equally high CD19-CAR expression rates (>85%; Figure 3b), high T-cell viability (>90%; Fig. 3b) and equally potent cytotoxicity against B-cells (Figure 3c). These results demonstrate that in applications of mRNA to T-cells, uridine modifications could be omitted in favour of unmodified mRNA.

**Figure 3.**
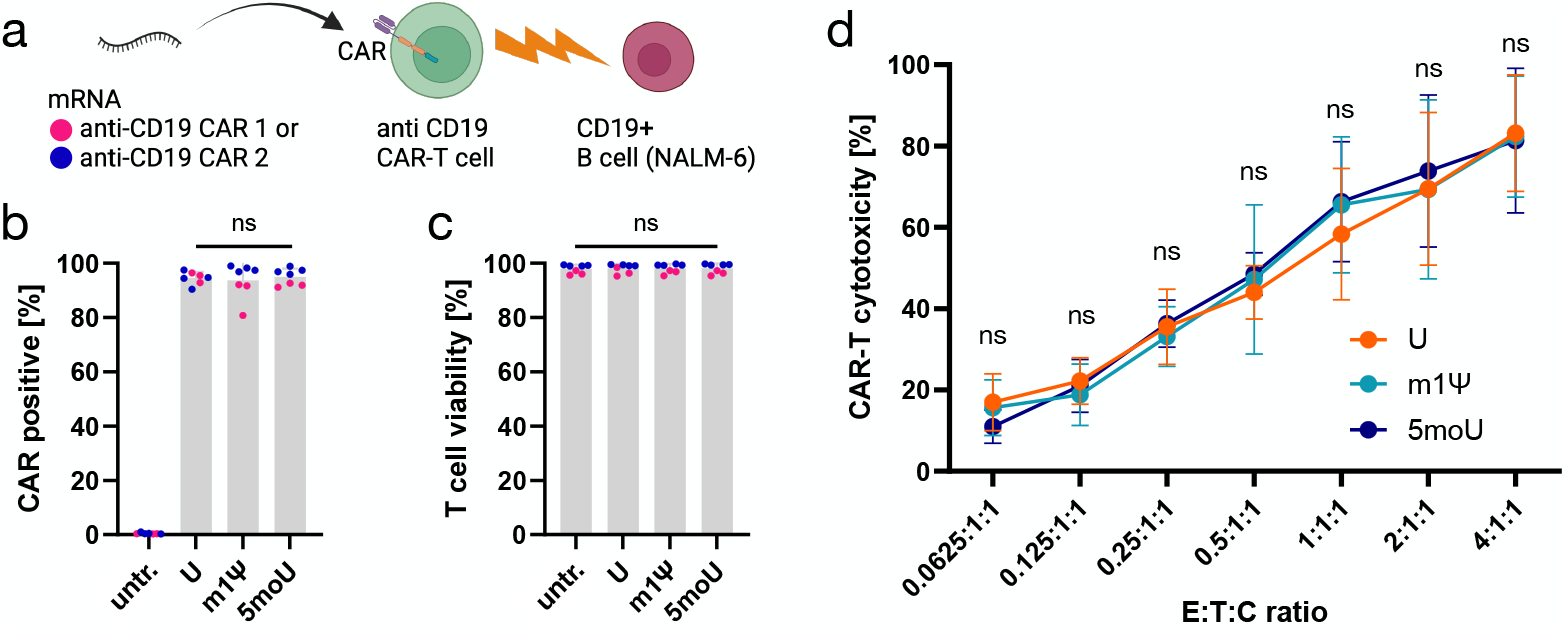
CAR-T cells generated with unmodified versus nucleotide-modified mRNA are functionally equivalent. **a)** Schematic: mRNA with or without nucleoside modifications encoding two different anti-CD19 chimeric antigen receptors (designated by magenta and blue respectively) was used to generate CAR-T cells directed against a target B cell line (NALM-6 cells). > 85% of T-cells showed CAR expression **(b)** and high viability **(c)**, irrespective of the mRNA nucleoside modification used. n= 7 donors. **d)** A VITAL assay to quantify antigen-specific cytotoxicity of CAR-T cells against NALM-6 B cells [16] was performed at an increasing ratio (E:T:C) between effector CAR-T cells (E), CD19+ target cells (T) and CD19-control cells (C). It revealed no difference in killing functionality between CAR-T cells generated with unmodified versus m1Ψ-modified or 5moU-modified mRNA. n=6: n=3 donors and n=2 CAR constructs. ns= not significant.

Compared to virus-transduction of CAR-T-cells, mRNA-based manufacturing could reduce cost, time and complexity[5]. Nevertheless, mRNA is commonly limited by antiviral immune responses. Uridine modification used to mitigate this immune activation account for ∼35% of IVT-mRNA manufacturing costs. However, human T-cells appear to lack receptors required to recognize uridine on exogenous mRNA. This study demonstrates that unmodified mRNA does not elicit immune reactions in T-cells, highlighting it as a safe, effective, and cost-efficient alternative for CAR-T cell manufacturing.

## Acknowledgements

This project had received funding from the Einstein Center for Regenerative Therapies (ECRT) Advanced Scientist Kickbox. Funding was provided by the Helmholtz Association and by the Federal Ministry of Education and Research, Germany, in the Program Health Research (BCRT grant no. 13GW0098 and 13GW0099). The authors would like to thank Lisa Hemmerling and Yusuf Aydemir for their help with experiments. The authors also thank Prof. Nina Babel (BCRT) for providing renal TEC cells and the BCRT Cell Harvesting Core Unit (BCRT-CH) of the Berlin Institute of Health, Charité – Universitätsmedizin Berlin for their excellent technical assistance and support. Figures were created using BioRender.com and GraphPad Prism.

## Authorship contributions

NoKa: Investigation, Formal analysis, Visualization. NiKo: Investigation, Formal analysis. JMP: Investigation, Visualization. YP: Investigation, Formal analysis. JK: Validation, Methodology. SP: Methodology. ALH: Software. CMD: Methodology. ML: Methodology, Writing – review & editing. AK: Writing – review & editing. DNS: Writing – review & editing. DLW: Methodology, Resources. CP: Writing – review & editing. GK: Writing – review & editing. MSH: Methodology, Resources, Funding acquisition, Writing – review & editing. HDV: Conceptualization, Funding acquisition, Writing – review & editing. MG: Conceptualization, Supervision, Resources, Writing – review & editing. NMD: Conceptualization, Project administration, Supervision, Methodology, Investigation, Formal analysis, Visualization, Writing – original draft, Funding acquisition.

## Data availability

All underlying data of this manuscript can be obtained from the corresponding author upon request. The pRNA2-(A)128 vector is available on Addgene (ID: 174006). The plasmid containing the CD19-CAR 2 sequence can be obtained from Addgene (ID: 183473).

## References

1. Hernandez, I., V. Prasad, and W.F. Gellad, Total Costs of Chimeric Antigen Receptor T-Cell Immunotherapy. JAMA Oncol, 2018. 4(7): p. 994–996.

2. Muller, F., J. Taubmann, L. Bucci, A. Wilhelm, C. Bergmann, S. Volkl, M. Aigner, T. Rothe, I. Minopoulou, C. Tur, J. Knitza, S. Kharboutli, S. Kretschmann, I. Vasova, S. Spoerl, H. Reimann, L. Munoz, R.G. Gerlach, S. Schafer, R. Grieshaber-Bouyer, A.S. Korganow, D. Farge-Bancel, D. Mougiakakos, A. Bozec, T. Winkler, G. Kronke, A. Mackensen, and G. Schett, CD19 CAR T-Cell Therapy in Autoimmune Disease - A Case Series with Follow-up. N Engl J Med, 2024. 390(8): p. 687–700.

3. Aghajanian, H., T. Kimura, J.G. Rurik, A.S. Hancock, M.S. Leibowitz, L. Li, J. Scholler, J. Monslow, A. Lo, W. Han, T. Wang, K. Bedi, M.P. Morley, R.A. Linares Saldana, N.A. Bolar, K. McDaid, C.A. Assenmacher, C.L. Smith, D. Wirth, C.H. June, K.B. Margulies, R. Jain, E. Pure, S.M. Albelda, and J.A. Epstein, Targeting cardiac fibrosis with engineered T cells. Nature, 2019. 573(7774): p. 430–433.

4. Amor, C., J. Feucht, J. Leibold, Y.J. Ho, C. Zhu, D. Alonso-Curbelo, J. Mansilla-Soto, J.A. Boyer, X. Li, T. Giavridis, A. Kulick, S. Houlihan, E. Peerschke, S.L. Friedman, V. Ponomarev, A. Piersigilli, M. Sadelain, and S.W. Lowe, Senolytic CAR T cells reverse senescence-associated pathologies. Nature, 2020. 583(7814): p. 127–132.

5. Baker, D.J., Z. Arany, J.A. Baur, J.A. Epstein, and C.H. June, CAR T therapy beyond cancer: the evolution of a living drug. Nature, 2023. 619(7971): p. 707–715.

6. Granit, V., M. Benatar, M. Kurtoglu, M.D. Miljkovic, N. Chahin, G. Sahagian, M.H. Feinberg, A. Slansky, T. Vu, C.M. Jewell, M.S. Singer, M.V. Kalayoglu, J.F. Howard, Jr., T. Mozaffar, and M.G.S. Team, Safety and clinical activity of autologous RNA chimeric antigen receptor T-cell therapy in myasthenia gravis (MG-001): a prospective, multicentre, open-label, non-randomised phase 1b/2a study. Lancet Neurol, 2023. 22(7): p. 578–590.

7. Meister, H., T. Look, P. Roth, S. Pascolo, U. Sahin, S. Lee, B.D. Hale, B. Snijder, L. Regli, V.M. Ravi, D.H. Heiland, C.L. Sentman, M. Weller, and T. Weiss, Multifunctional mRNA-Based CAR T Cells Display Promising Antitumor Activity Against Glioblastoma. Clin Cancer Res, 2022. 28(21): p. 4747–4756.

8. Sahin, U., K. Kariko, and O. Tureci, mRNA-based therapeutics--developing a new class of drugs. Nat Rev Drug Discov, 2014. 13(10): p. 759–80.

9. Nance, K.D. and J.L. Meier, Modifications in an Emergency: The Role of N1-Methylpseudouridine in COVID-19 Vaccines. ACS Cent Sci, 2021. 7(5): p. 748–756.

10. Drzeniek, N.M., N. Kahwaji, S. Schlickeiser, P. Reinke, S. Geißler, H.-D. Volk, and M. Gossen, Immuno-engineered mRNA combined with cell adhesive niche for synergistic modulation of the MSC secretome. Biomaterials, 2023. 294: p. 121971.

11. Drzeniek, N.M., N. Kahwaji, S. Picht, I.M. Dimitriou, S. Schlickeiser, H. Moradian, S. Geissler, M. Schmueck-Henneresse, M. Gossen, and H.D. Volk, In Vitro Transcribed mRNA Immunogenicity Induces Chemokine-Mediated Lymphocyte Recruitment and Can Be Gradually Tailored by Uridine Modification. Adv Sci (Weinh), 2024: p. e2308447.

12. Billingsley, M.M., N. Gong, A.J. Mukalel, A.S. Thatte, R. El-Mayta, S.K. Patel, A.E. Metzloff, K.L. Swingle, X. Han, L. Xue, A.G. Hamilton, H.C. Safford, M.G. Alameh, T.E. Papp, H. Parhiz, D. Weissman, and M.J. Mitchell, In Vivo mRNA CAR T Cell Engineering via Targeted Ionizable Lipid Nanoparticles with Extrahepatic Tropism. Small, 2024. 20(11): p. e2304378.

13. Kitte, R., M. Rabel, R. Geczy, S. Park, S. Fricke, U. Koehl, and U.S. Tretbar, Lipid nanoparticles outperform electroporation in mRNA-based CAR T cell engineering. Mol Ther Methods Clin Dev, 2023. 31: p. 101139.

14. Moradian, H., T. Roch, L. Anthofer, A. Lendlein, and M. Gossen, Chemical modification of uridine modulates mRNA-mediated proinflammatory and antiviral response in primary human macrophages. Mol Ther Nucleic Acids, 2022. 27: p. 854–869.

15. Drzeniek, N.M., A. Mazzocchi, S. Schlickeiser, S.D. Forsythe, G. Moll, S. Geissler, P. Reinke, M. Gossen, V.S. Gorantla, H.D. Volk, and S. Soker, Bio-instructive hydrogel expands the paracrine potency of mesenchymal stem cells. Biofabrication, 2021. 13(4).

16. Kath, J., C. Franke, V. Drosdek, W.J. Du, V. Glaser, C. Fuster-Garcia, M. Stein, T. Zittel, S. Schulenberg, C.E. Porter, L. Andersch, A. Künkele, J. Alcaniz, J. Hoffmann, H. Abken, M. Abou-el-Enein, A. Pruss, M. Suzuki, T. Cathomen, R. Stripecke, H.D. Volk, P. Reinke, M. Schmueck-Henneresse, and D.L. Wagner, Integration of ζ-deficient CARs into the CD3ζ gene conveys potent cytotoxicity in T and NK cells. Blood, 2024. 143(25): p. 2599–2611.

17. Kath, J., W. Du, A. Pruene, T. Braun, B. Thommandru, R. Turk, M.L. Sturgeon, G.L. Kurgan, L. Amini, M. Stein, T. Zittel, S. Martini, L. Ostendorf, A. Wilhelm, L. Akyuz, A. Rehm, U.E. Hopken, A. Pruss, A. Kunkele, A.M. Jacobi, H.D. Volk, M. Schmueck-Henneresse, R. Stripecke, P. Reinke, and D.L. Wagner, Pharmacological interventions enhance virus-free generation of TRAC-replaced CAR T cells. Mol Ther Methods Clin Dev, 2022. 25: p. 311–330.

18. Hammoud, B., M. Schmueck, A.M. Fischer, H. Fuehrer, S.J. Park, L. Akyuez, J.C. Schefold, M.J. Raftery, G. Schonrich, A.M. Kaufmann, H.D. Volk, and P. Reinke, HCMV-specific T-cell therapy: do not forget supply of help. J Immunother, 2013. 36(2): p. 93–101.

19. Monaco, G., B. Lee, W. Xu, S. Mustafah, Y.Y. Hwang, C. Carre, N. Burdin, L. Visan, M. Ceccarelli, M. Poidinger, A. Zippelius, J. Pedro de Magalhaes, and A. Larbi, RNA-Seq Signatures Normalized by mRNA Abundance Allow Absolute Deconvolution of Human Immune Cell Types. Cell Rep, 2019. 26(6): p. 1627–1640 e7.

20. Schmiedel, B.J., D. Singh, A. Madrigal, A.G. Valdovino-Gonzalez, B.M. White, J. Zapardiel-Gonzalo, B. Ha, G. Altay, J.A. Greenbaum, G. McVicker, G. Seumois, A. Rao, M. Kronenberg, B. Peters, and P. Vijayanand, Impact of Genetic Polymorphisms on Human Immune Cell Gene Expression. Cell, 2018. 175(6): p. 1701–1715 e16.

