## Supplemental Information for "Revisiting the Need for mRNA Nucleoside Modification in CAR T Cell Engineering"

**List of Supplemental Information:**

- Supplemental Table 1: Open reading frames of EGFP and CARs used in this study.
- Figure S1. Lipofection of primary human T cells using lipofectamine complexes versus lipid nanoparticles
- Figure S2. Representative gating strategy for flow cytometry.
- Figure S3. Interferon production in response to mRNA with different uridine chemistry varies between cell types.
- Figure S4. Cell death in response to mRNA with different uridine chemistry varies between cell types.
- Figure S5. Expression of mRNA with different uridine chemistry varies between cell types.
- Figure S6. T cells from four donors transfected with unmodified mRNA.
- Figure S7. Expression of mRNA-recognizing receptors in human immune cell subsets.

**Supplemental Table 1:**

| Target name | coding sequence | Ref. |
| --- | --- | --- |
| eGFP | ATGGTGAGCAAGGGCGAGGAGCTGTTCACCGGGGTGGTGCCCATCCTGGTCGAGCTGGACGGCGACGTAAACGGCCACAAGTTCAGCGTGTCCGGCGAGGGCGAGGGCGATGCCACCTACGGCAAGCTGACCCTGAAGTTCATCTGCACCACCGGCAAGCTGCCCGTGCCCTGGCCCACCCTCGTGACCACCCTGACCTACGGCGTGCAGTGCTTCAGCCGCTACCCCGACCACATGAAGCAGCACGACTTCTTCAAGTCCGCCATGCCCGAAGGCTACGTCCAGGAGCGCACCATCTTCTTCAAGGACGACGGCAACTACAAGACCCGCGCCGAGGTGAAGTTCGAGGGCGACACCCTGGTGAACCGCATCGAGCTGAAGGGCATCGACTTCAAGGAGGACGGCAACATCCTGGGGCACAAGCTGGAGTACAACTACAACAGCCACAACGTCTATATCATGGCCGACAAGCAGAAGAACGGCATCAAGGTGAACTTCAAGATCCGCCACAACATCGAGGACGGCAGCGTGCAGCTCGCCGACCACTACCAGCAGAACACCCCCATCGGCGACGGCCCCGTGCTGCTGCCCGACAACCACTACCTGAGCACCCAGTCCGCCCTGAGCAAAGACCCCAACGAGAAGCGCGATCACATGGTCCTGCTGGAGTTCGTGACCGCCGCCGGGATCACTCTCGGCATGGACGAGCTGTACAAGTAA | [1] |
| CD19-CAR 1 (anti-Myc stainable) | ATGGCTCTTCCTGTGACTGCCCTTCTGCTGCCCCTGGCTCTCCTGCTCCATGCTGCCCGGCCAgagcaaaaacttatctctgaagaggacctcGACATCCAGATGACACAGACAACCAGCAGCCTCAGCGCCAGCCTGGGGGACAGAGTGACCATTAGCTGCCGGGCCTCTCAGGACATCAGCAAATACCTGAACTGGTACCAGCAGAAACCAGATGGCACTGTCAAGCTGCTGATTTACCACACCTCCAGGCTCCACAGCGGCGTGCCCAGTCGCTTCAGCGGCAGTGGGAGCGGGACAGATTATTCCCTCACAATCTCCAACCTGGAGCAGGAAGATATTGCCACATACTTCTGCCAGCAAGGCAACACCCTGCCATACACATTTGGAGGCGGCACCAAATTGGAGATCACCGGCGGTGGTGGATCTGGAGGAGGGGGCAGCGGAGGTGGCGGCTCTGAGGTGAAACTGCAGGAGAGTGGCCCTGGCCTGGTGGCTCCCAGCCAGAGCCTTTCTGTCACCTGCACCGTGTCTGGGGTGTCCCTGCCTGACTATGGAGTCTCCTGGATCCGGCAGCCTCCAAGAAAAGGACTGGAATGGCTGGGCGTCATCTGGGGAAGTGAGACCACCTACTATAATTCAGCCCTCAAGTCCCGGCTCACCATCATTAAGGACAACTCCAAATCCCAGGTGTTCCTGAAGATGAATTCTCTCCAGACTGATGACACAGCCATCTACTACTGTGCCAAGCATTATTATTACGGCGGGTCCTATGCCATGGACTACTGGGGCCAGGGGACCAGTGTCACTGTTTCTTCTATAGAAGTAATGTATCCCCCTCCCTACTTGGACAACGAGAAATCTAACGGCACAATCATACACGTTAAGGGCAAGCATCTGTGTCCCTCCCCTCTTTTCCCCGGACCGTCTAAGCCATTTTGGGTCCTCGTGGTGGTCGGGGGTGTGCTGGCCTGCTACAGCTTGCTGGTCACAGTGGCCTTCATCATCTTCTGGGTGCGCTCCAAGAGGAGCCGGCTGCTTCACAGTGATTACATGAACATGACCCCCAGGAGGCCAGGACCCACCAGGAAGCACTACCAGCCCTACGCTCCCCCGCGGGACTTTGCTGCTTACCGCAGCAGGGTCAAATTTTCTAGATCTGCAGATGCGCCGGCCTATCAAcaaggccagaaccagctcTATAACGAGCTCAATCTAGGACGAAGAGAGGAGTACGATGTTTTGGACAAGAGACGTGGCCGGGACCCTGAGATGGGGGGAAAGCCGAGAAGGAAGAACCCTCAGGAAGGCCTGTACAATGAACTGCAGAAAGATAAGATGGCGGAGGCCTACAGTGAGATTGGGATGAAAGGCGAGCGCCGGAGGGGCAAGGGGCACGATGGCCTTTACCAGGGTCTCAGTACAGCCACCAAGGACACCTACGACGCCCTTCACATGCAGGCCCTGCCCCCTCGCTAA | [2] |
| CD19-CAR 2  (anti-Fc stainable) | ATGGCTCTTCCTGTGACTGCCCTTCTGCTGCCCCTGGCTCTCCTGCTCCATGCTGCCCGGCCAGACATCCAGATGACACAGACAACCAGCAGCCTCAGCGCCAGCCTGGGGGACAGAGTGACCATTAGCTGCCGGGCCTCTCAGGACATCAGCAAATACCTGAACTGGTACCAGCAGAAACCAGATGGCACTGTCAAGCTGCTGATTTACCACACCTCCAGGCTCCACAGCGGCGTGCCCAGTCGCTTCAGCGGCAGTGGGAGCGGGACAGATTATTCCCTCACAATCTCCAACCTGGAGCAGGAAGATATTGCCACATACTTCTGCCAGCAAGGCAACACCCTGCCATACACATTTGGAGGCGGCACCAAATTGGAGATCACCGGCGGTGGTGGATCTGGAGGAGGGGGCAGCGGAGGTGGCGGCTCTGAGGTGAAACTGCAGGAGAGTGGCCCTGGCCTGGTGGCTCCCAGCCAGAGCCTTTCTGTCACCTGCACCGTGTCTGGGGTGTCCCTGCCTGACTATGGAGTCTCCTGGATCCGGCAGCCTCCAAGAAAAGGACTGGAATGGCTGGGCGTCATCTGGGGAAGTGAGACCACCTACTATAATTCAGCCCTCAAGTCCCGGCTCACCATCATTAAGGACAACTCCAAATCCCAGGTGTTCCTGAAGATGAATTCTCTCCAGACTGATGACACAGCCATCTACTACTGTGCCAAGCATTATTATTACGGCGGGTCCTATGCCATGGACTACTGGGGCCAGGGGACCAGTGTCACTGTTTCTTCTGAAAGCAAGTATGGCCCCCCATGCCCTCCCTGCCCCGGACAGCCGCGGGAGCCCCAGGTCTACACCCTCCCACCCTCCAGAGATGAGCTGACCAAGAACCAGGTTTCACTGACATGCCTGGTGAAGGGCTTCTACCCCTCTGACATTGCTGTGGAGTGGGAGAGCAATGGGCAGCCAGAGAACAACTACAAGACCACACCTCCTGTCCTGGACAGCGACGGCTCCTTCTTCCTCTACAGTAAGCTCACTGTGGACAAGAGCCGCTGGCAGCAGGGCAATGTCTTCTCCTGCAGCGTGATGCACGAGGCCCTGCACAATGCCTATACCCAGAAGTCACTCTCTCTGAGCCCTGGAAAGAAGGATCCCAAGTTTTGGGTCCTCGTGGTGGTCGGGGGTGTGCTGGCCTGCTACAGCTTGCTGGTCACAGTGGCCTTCATCATCTTCTGGGTGCGCTCCAAGAGGAGCCGGCTGCTTCACAGTGATTACATGAACATGACCCCCAGGAGGCCAGGACCCACCAGGAAGCACTACCAGCCCTACGCTCCCCCGCGGGACTTTGCTGCTTACCGCAGCAGGGTCAAATTTTCTAGATCTGCAGATGCGCCGGCCTATCAAcaaggccagaaccagctcTATAACGAGCTCAATCTAGGACGAAGAGAGGAGTACGATGTTTTGGACAAGAGACGTGGCCGGGACCCTGAGATGGGGGGAAAGCCGAGAAGGAAGAACCCTCAGGAAGGCCTGTACAATGAACTGCAGAAAGATAAGATGGCGGAGGCCTACAGTGAGATTGGGATGAAAGGCGAGCGCCGGAGGGGCAAGGGGCACGATGGCCTTTACCAGGGTCTCAGTACAGCCACCAAGGACACCTACGACGCCCTTCACATGCAGGCCCTGCCCCCTCGCTAA | [3] |

**Supplemental Table 1. Open reading frames of EGFP and CARs used in this study.**

A reference for each sequence is provided in the right column.

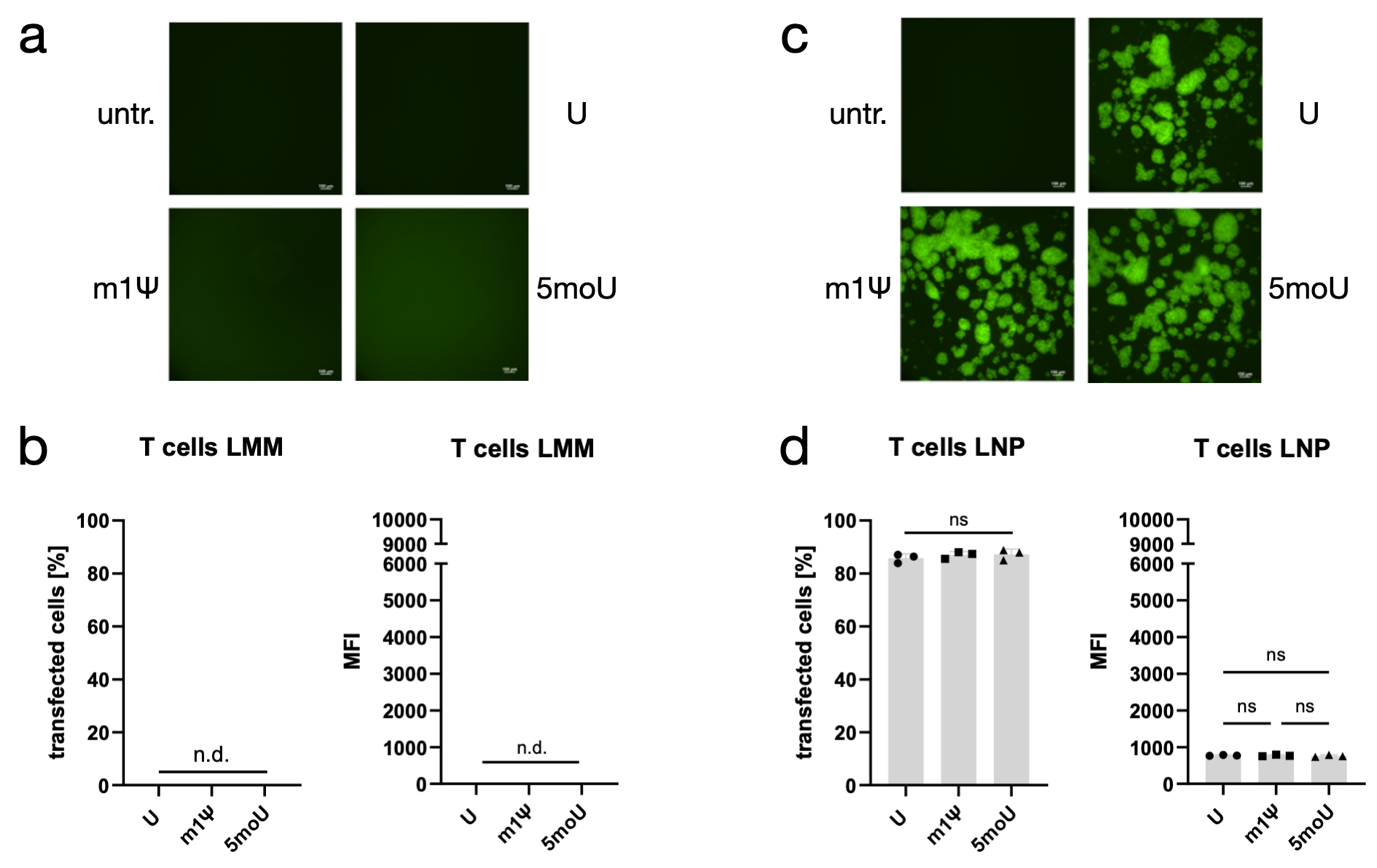

**Supplemental Figure S1. Lipofection of primary human T cells using lipofectamine complexes versus lipid nanoparticles.**  Primary human T cells isolated from the blood of 3 healthy donors were activated for 2 days and treated with with unmodified (U) or m1Ψ-modified or 5moU-modified GFP-encoding mRNA, complexed with a lipid delivery vehicle. Complexation of mRNA with the transfection reagent Lipofectamine MessengerMax (LMM) did not reasult in any T cells transfection, as observed **(a)** using a Nikon ECLIPSE Ti fluorescence microscope (Nikon Instruments Inc., Japan) or quantified by flow cytometry (MACSQuant VYB, Miltenyi Biotec, Germany) **(b)**. In contrast, mRNA complexed with Genvoy lipid mix did result in efficient T cell transfection **(c, d)**. d) shows the same data as Figure S5e, j. Microscopy settings were kept identical between images. GFP fluorescence is shown in green. n = 3. * p<0,05; ** p<0,01; *** p<0,001; ns= not significant.

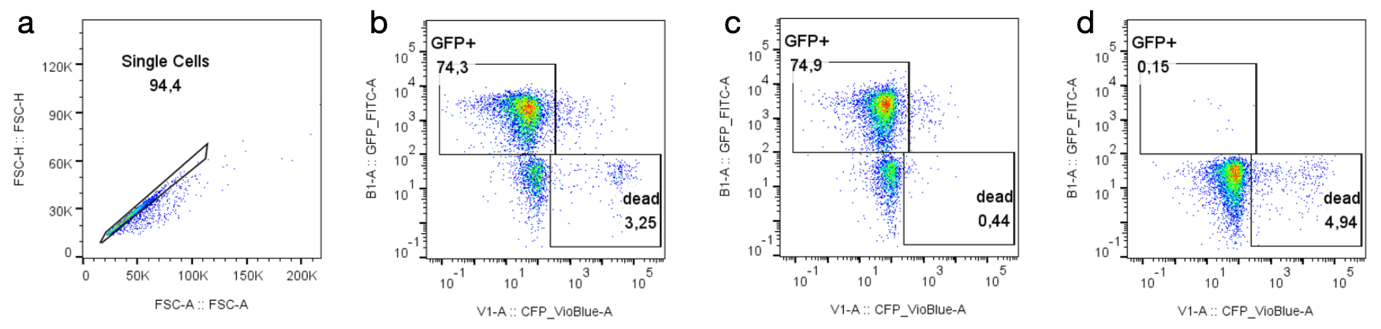

**Supplemental Figure S2. Representative gating strategy for flow cytometry.** The gating strategy is shown for a purified population of T cells but applies to all cell types in this study. After defining the cell population in the FSC/SSC plot (not shown), single cells were selected using FSC height against area **(a)**. Next, the channels detecting GFP (B1) and DAPI (V1) were plotted against each other, in order to determine GFP and DAPI positive cells **(b)**. Here a sample of T cells electroporated with GFP-encoding mRNA is shown. The same sample was measured before adding DAPI to set the gate for live/dead discrimination **(c)**. A control sample, which was not treated with GFP-mRNA was used to set the GFP gate **(d)**. Data was aquired using a MACSQuant VYB cytometer (Miltenyi Biotec, Germany). Data analysis was carried out using FlowJo v10 (BD Biosciences, USA).

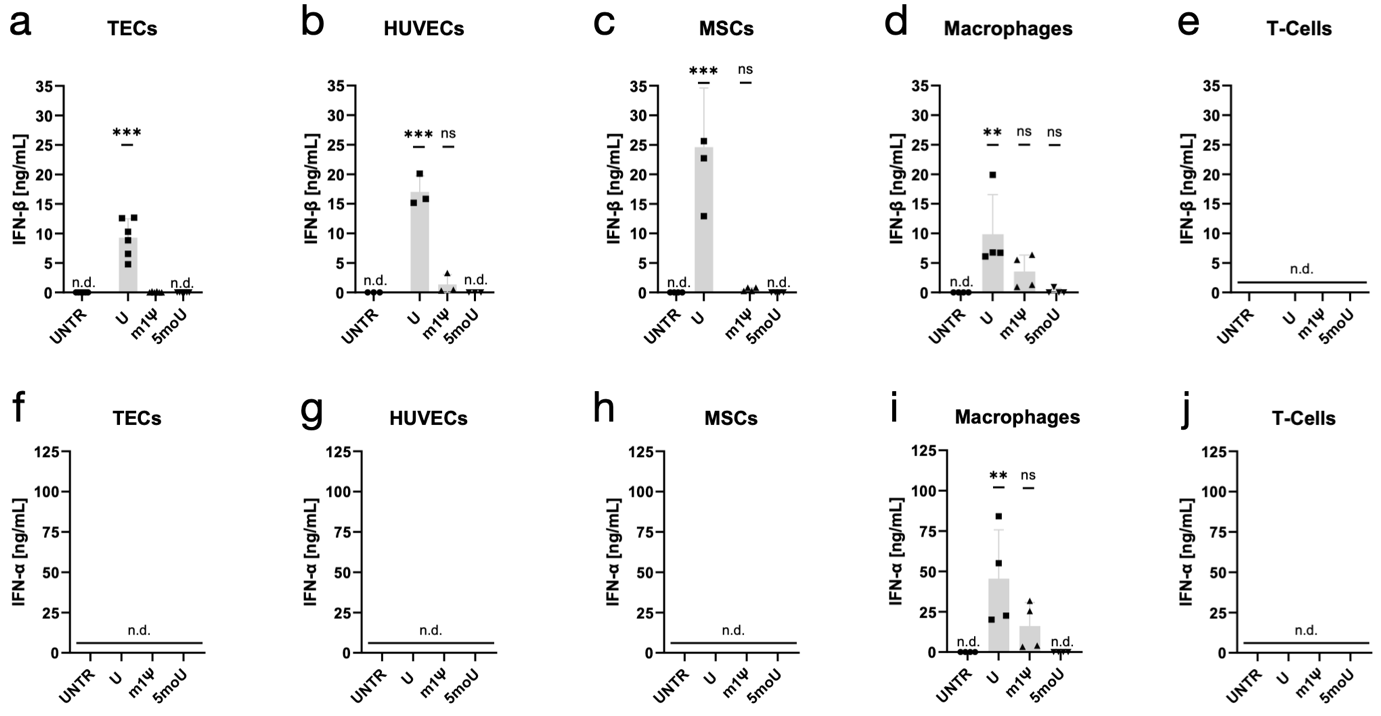

**Supplemental Figure S3. Interferon production in response to mRNA with different uridine chemistry varies between cell types.**   Renal tubular epithelial cells (TECs), umbilical vein endothelial cells (HUVECs), bone marrow stromal cells (MSCs), macrophages and T cells were transfected with unmodified (U) or m1Ψ-modified or 5moU-modified GFP-encoding mRNA and concentration of type-I interferons β (a-e) and α (f-j) in supernatants was measured 24h post-transfection. n ≥ 3. * p<0,05; ** p<0,01; *** p<0,001; ns= not significant.

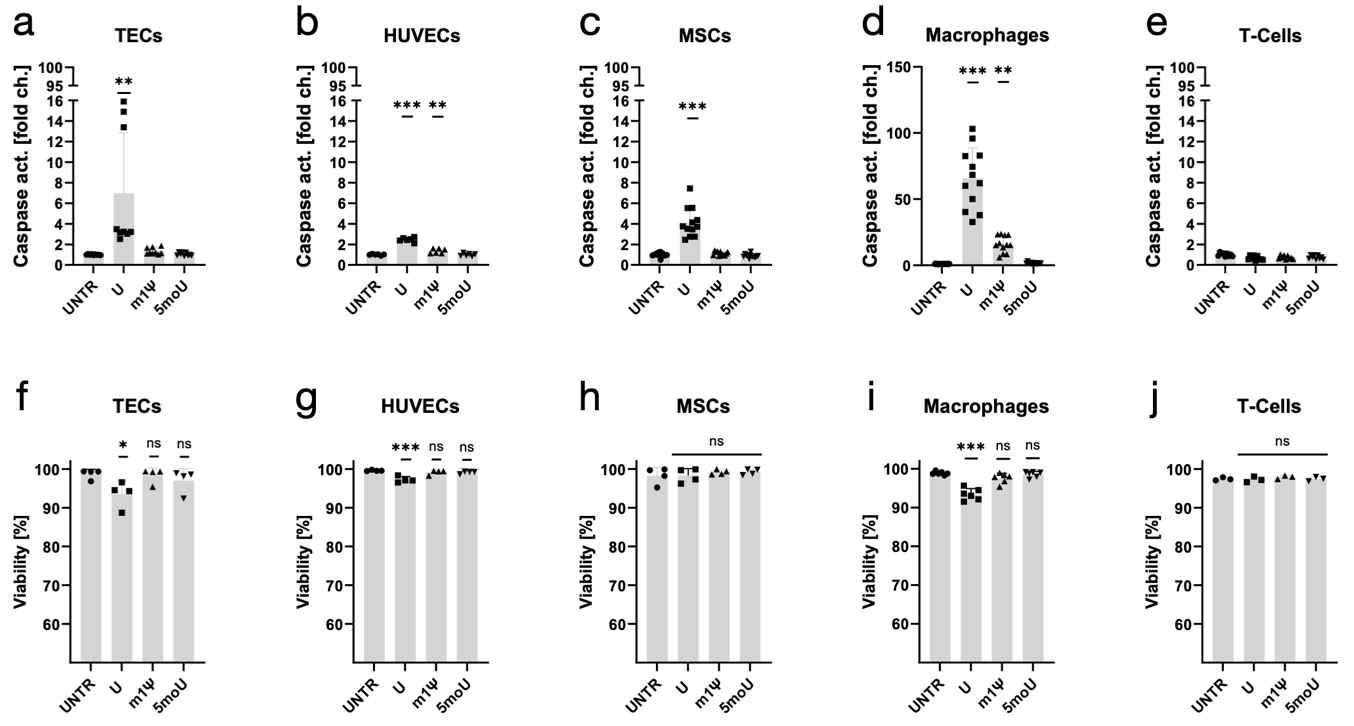
**Supplemental Figure S4. Cell death in response to mRNA with different uridine chemistry varies between cell types.**   Renal tubular epithelial cells (TECs), umbilical vein endothelial cells (HUVECs), bone marrow stromal cells (MSCs), macrophages and T cells were transfected with unmodified (U) or m1Ψ-modified or 5moU-modified GFP-encoding mRNA. Increase in caspase activity, expressed as fold change from untransfected cells, was measured using a luminescence-based assay for caspase 3 and 7 activity (a-e). Cell death was quantified 24h post-transfection using DAPI staining and flow cytometry (f-j). n ≥ 3.* p<0,05; ** p<0,01; *** p<0,001; ns= not significant.

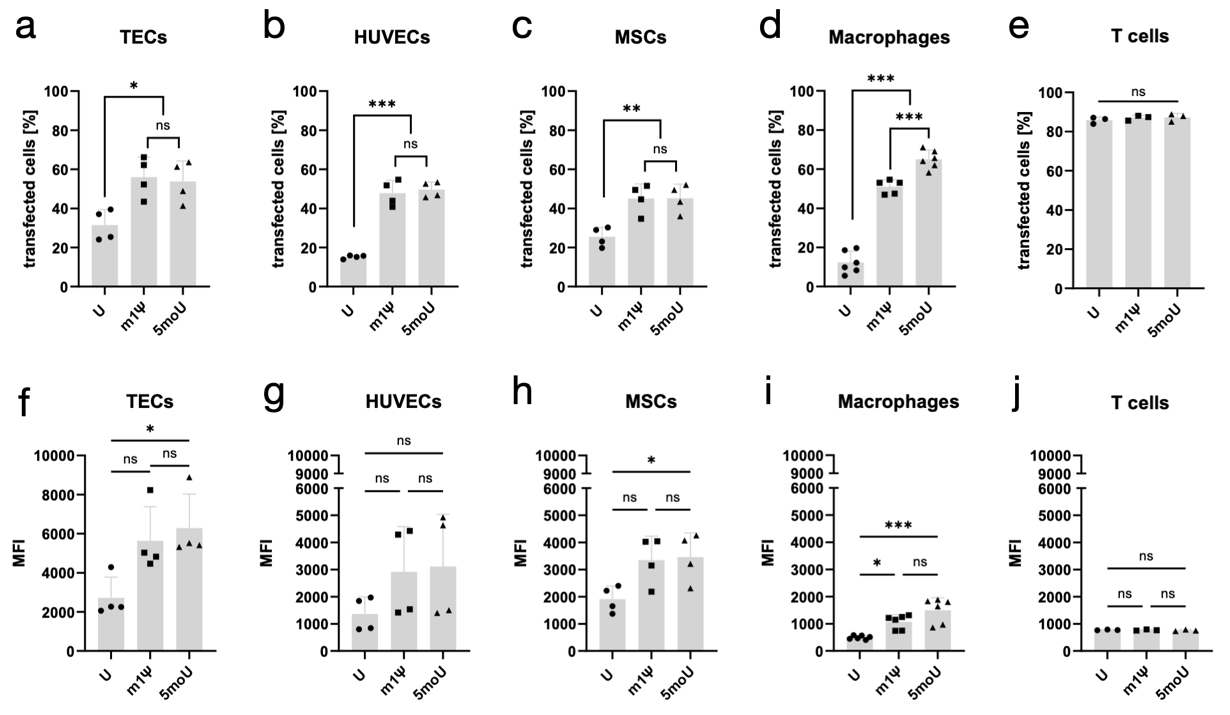
**Supplemental Figure S5. Expression of mRNA with different uridine chemistry varies between cell types.**   Renal tubular epithelial cells (TECs), umbilical vein endothelial cells (HUVECs), bone marrow stromal cells (MSCs), macrophages and T cells were transfected with unmodified (U) or m1Ψ-modified or 5moU-modified GFP-encoding mRNA. The percentage of GFP positive cells (a-e) and the mean fluorescence intensity (MFI) of the GFP fluorescence (f-j) were quantified using flow cytometry with equal GFP-laser setting for all measurements. n ≥ 3.* p<0,05; ** p<0,01; *** p<0,001; ns= not significant.

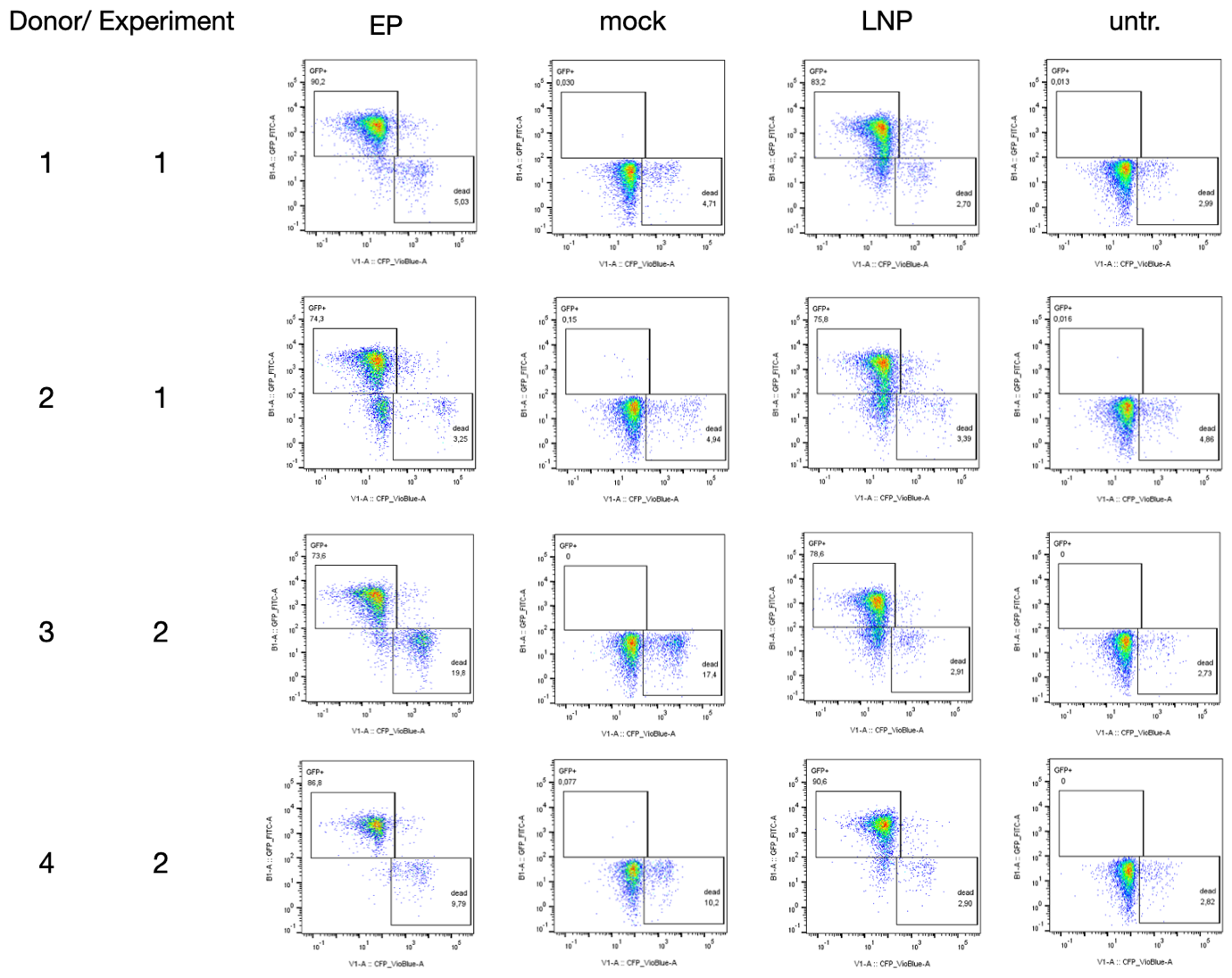

**Supplemental Figure S6. T cells from four donors transfected with unmodified mRNA.** Primary human T cells isolated from the blood of 4 healthy donors were activated for 2 days and transfected with unmodified GFP-encoding mRNA either by electroporation (EP) or lipid nanoparticle (LNP) delivery in 2 separate experiments. Untreated cells (untr.) and cells subjected to the electroporation procedure but without mRNA (mock) served as controls. n=1 out of 3 technical replicates is shown for each donor. Across donors and experiments, unmodified mRNA did not reduce T cell viability and yielded high expression, irrespective of the transfection method. Data was aquired using a MACSQuant VYB cytometer (Miltenyi Biotec, Germany). Data analysis was carried out using FlowJo v10 (BD Biosciences, USA).

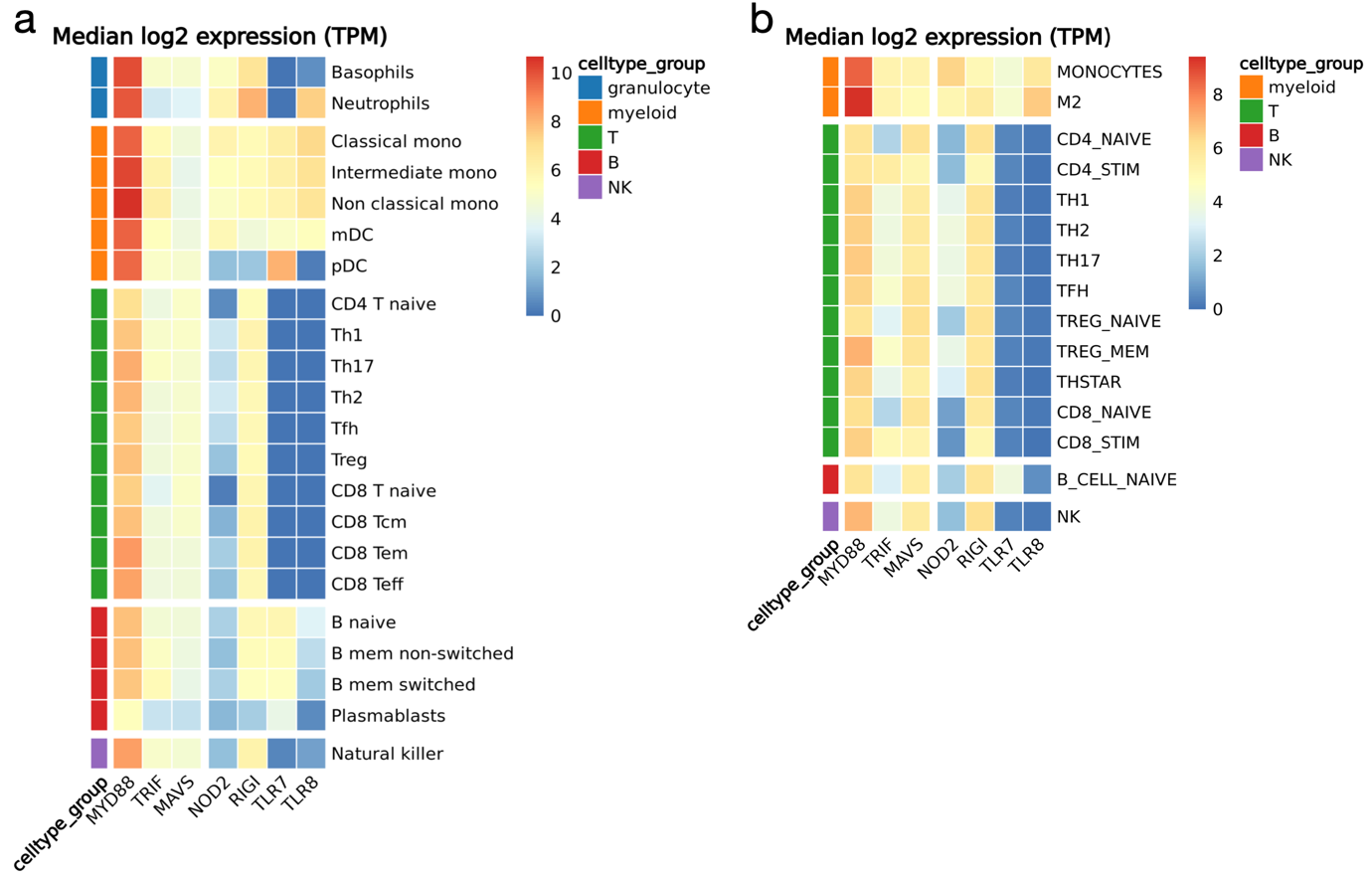
**Supplemental** **Figure S7. Expression of mRNA-recognizing receptors in human immune cell subsets.**   Two independent, published and publicly available datasets of bulk RNA sequencing data were interrogated for RNA expression levels of pattern recognition receptors (PRR; TLR7, TLR8, RIGI, NOD2) against exogenous single stranded mRNA and adapter proteins (MyD88, TRIF and MAVS). Among those PRR, TLR 7 and 8 recognize unmodified uridine on single-stranded mRNA[4, 5]. Both datasets show that these two receptors were not expressed in any T lymphocyte subsets and their adapters MyD88 and TRIF were expressed at lower levels in T cells than in other leukocytes. Data wrangling and visualization was done in R v4.4.2. **(a)** RNA expression data from the Monaco et al. dataset [6] was accessed via GEO accession number GSE107011 using the GEOquery package (version 2.74.0). Only a subset of cell types was selected. (a) is also shown in the main article as Figure 2e. **(b)** Data referring to Schmiedel et al. [7] was downloaded from the DICE (Database of Immune Cell Expression, Expression quantitative trait loci (eQTLs) and Epigenomics) database (accessed on October 2, 2024). RNA expression data for each cell type, reported as transcripts per million (TPM), was used. For both datasets, median TPM values per cell type were calculated and log2(x + 1) transformed using dplyr v1.1.4 and tidyr v1.3.1. Data was visualized using the pheatmap package v1.0.12.

1.	Drzeniek, N.M., N. Kahwaji, S. Picht, I.M. Dimitriou, S. Schlickeiser, H. Moradian, S. Geissler, M. Schmueck-Henneresse, M. Gossen, and H.D. Volk, *In Vitro Transcribed mRNA Immunogenicity Induces Chemokine-Mediated Lymphocyte Recruitment and Can Be Gradually Tailored by Uridine Modification.* Adv Sci (Weinh), 2024: p. e2308447.

2.	Kath, J., C. Franke, V. Drosdek, W.J. Du, V. Glaser, C. Fuster-Garcia, M. Stein, T. Zittel, S. Schulenberg, C.E. Porter, L. Andersch, A. Künkele, J. Alcaniz, J. Hoffmann, H. Abken, M. Abou-el-Enein, A. Pruss, M. Suzuki, T. Cathomen, R. Stripecke, H.D. Volk, P. Reinke, M. Schmueck-Henneresse, and D.L. Wagner, *Integration of ζ-deficient CARs into the CD3ζ gene conveys potent cytotoxicity in T and NK cells.* Blood, 2024. **143**(25): p. 2599-2611.

3.	Kath, J., W. Du, A. Pruene, T. Braun, B. Thommandru, R. Turk, M.L. Sturgeon, G.L. Kurgan, L. Amini, M. Stein, T. Zittel, S. Martini, L. Ostendorf, A. Wilhelm, L. Akyuz, A. Rehm, U.E. Hopken, A. Pruss, A. Kunkele, A.M. Jacobi, H.D. Volk, M. Schmueck-Henneresse, R. Stripecke, P. Reinke, and D.L. Wagner, *Pharmacological interventions enhance virus-free generation of TRAC-replaced CAR T cells.* Mol Ther Methods Clin Dev, 2022. **25**: p. 311-330.

4.	Karikó, K., M. Buckstein, H. Ni, and D. Weissman, *Suppression of RNA Recognition by Toll-like Receptors: The Impact of Nucleoside Modification and the Evolutionary Origin of RNA.* Immunity, 2005. **23**(2): p. 165-175.

5.	Diebold, S.S., C. Massacrier, S. Akira, C. Paturel, Y. Morel, and C. Reis e Sousa, *Nucleic acid agonists for Toll-like receptor 7 are defined by the presence of uridine ribonucleotides.* Eur J Immunol, 2006. **36**(12): p. 3256-67.

6.	Monaco, G., B. Lee, W. Xu, S. Mustafah, Y.Y. Hwang, C. Carre, N. Burdin, L. Visan, M. Ceccarelli, M. Poidinger, A. Zippelius, J. Pedro de Magalhaes, and A. Larbi, *RNA-Seq Signatures Normalized by mRNA Abundance Allow Absolute Deconvolution of Human Immune Cell Types.* Cell Rep, 2019. **26**(6): p. 1627-1640 e7.

7.	Schmiedel, B.J., D. Singh, A. Madrigal, A.G. Valdovino-Gonzalez, B.M. White, J. Zapardiel-Gonzalo, B. Ha, G. Altay, J.A. Greenbaum, G. McVicker, G. Seumois, A. Rao, M. Kronenberg, B. Peters, and P. Vijayanand, *Impact of Genetic Polymorphisms on Human Immune Cell Gene Expression.* Cell, 2018. **175**(6): p. 1701-1715 e16.
